# Cortex-wide characterization of decision-making neural dynamics during spatial navigation

**DOI:** 10.1101/2024.10.23.619896

**Authors:** Samuel P. Haley, Daniel A. Surinach, Angela K. Nietz, Russell E. Carter, Lucas S. Zecker, Laurentiu S. Popa, Suhasa B. Kodandaramaiah, Timothy J. Ebner

## Abstract

Decision-making during freely moving behaviors involves complex interactions among many cortical and subcortical regions. However, the spatiotemporal coordination across regions to generate a decision is less understood. Using a head-mounted widefield microscope, cortex-wide calcium dynamics were recorded in mice expressing GCaMP7f as they navigated an 8-maze using two paradigms. The first was an alternating pattern that required short term memory of the previous trial to make the correct decision and the second after a rule change to a fixed path in which rewards were delivered only on the left side. Identification of cortex-wide activation states revealed differences between the two paradigms. There was a higher probability for a visual/retrosplenial cortical state during the alternating paradigm and higher probability of a secondary motor and posterior parietal state during left-only. Three state sequences (motifs) illustrated both anterior and posterior activity propagations across the cortex. The anterior propagating motifs had the highest probability around the decision and posterior propagating motifs peaked following the decision. The latter, likely reflecting internal feedback to influence future actions, were more common in the left-only paradigm. Therefore, the probabilities and sequences of cortical states differ when working memory is required versus a fixed trajectory reward paradigm.

## Introduction

Decisions are generated through complex interactions among various neocortical regions. The perception-action cycle is a common model used to describe decision-making behavior as a hierarchical flow of activation from posterior sensory regions to anterior motor regions^1^. As an organism collects and processes sensory information about its environment, this information is passed to higher-order cortical regions to weigh decision outcomes and execute commands. While this perception-action cycle predominantly follows the classical dorsal stream of activation with an anterior (feedforward) flow of activation^2^, it has also been hypothesized that the system operates in the reverse direction to provide feedback and influence future decision-making behavior^1,3,4^. Although coordinated feedforward/feedback signals have been proposed and cerebral cortical connectivity supports this concept (for reviews see^4–6^), there is limited evidence for bidirectional activation across the cortex as most studies have focused on information flows between two regions. Visualizing and quantifying reciprocal patterns of communication among many brain regions simultaneously is fundamental to understanding how the cortex functions as a cohesive system to generate decisions and behavior.

Cortical lesions and optogenetic inactivation in animals show that decision accuracy requires several key cortical regions. The retrosplenial cortex (RSP), posterior parietal cortex (PPC), and secondary motor cortex (M2) are all critically involved in various decision and spatial navigation tasks^4,7–11^. Inhibition of these regions effects decision accuracy only for specific assays, suggesting alternative circuits are utilized depending on the rules of the task^12,13^. While lesion and inhibition studies have provided essential demonstrations of the importance of single regions, approaches are needed that simultaneously assess the contributions of multiple regions to decision-making in real-time. Mesoscopic optical imaging of cortical dynamics is one approach to capture the roles of diverse regions during decision tasks^7,14,15^.

While wide-field calcium imaging has been proven to be a powerful method to investigate neocortical dynamics at the mesoscale level^12–23^, most investigations leverage this technique in head-fixed subjects. Expanding wide-field imaging in freely moving mice would allow investigation of more natural and complex behaviors as neural activity during free, unrestrained behavior can differ substantially from head-fixed recordings^24–26^. Therefore, there is a need to examine cortical engagement in freely moving behaviors to capture naturally occurring spatiotemporal patterns of neural activation.

Taking advantage of a recently developed miniaturized head-mounted microscope (mini-mScope^27,28^), we recorded cerebral cortex-wide calcium activity of freely-moving mice during two different paradigms in an 8-maze decision task. The two paradigms required different behavioral strategies to isolate the cortical circuitry engaged during decision-making. One paradigm required memory recall of the previous decision to correctly make the next decision (alternating), while the other paradigm simply required a fixed navigation pattern keeping all other aspects of the task consistent (left-only). While several forms of analyses have been used for widefield calcium imaging^7,14^, we utilized a form of brain state analysis that reduces widefield calcium data into discrete “states” to characterize cortex-wide activity patterns and how they change during the task^20,28–31^. The probabilities of the state activations and two state transitions differ between the two paradigms. By quantifying motifs using sequences of 3 states, we found not only anterior propagating motifs of cortical activation states consistent with the dorsal stream but also posterior propagating motifs. Differential utilizations of these feedforward and feedback spatiotemporal flows during the decision support the perception-action cycle and reveal a framework for the interactions among cortical regions underlying decision-making during spatial navigation.

## Results

### Wide-field imaging in freely moving mice and 8-maze paradigms

Mesoscale calcium imaging in mice expressing GCaMP7f using the head-mounted mini-mScope (**Figure 1A**) produces clear, high-quality images of a large region of the dorsal neocortex (**Figure 1B**). Post-mortem histological GFP labeling reveals dense, even expression of neurons throughout the cortex for both anterior (**Figure 1C**) and posterior (**Figure 1D**) cortical regions. Six mice were trained in a modified T-maze we will refer to as the 8-maze (**Figure 1E**). An automatic swing door was used to isolate a single decision at the top of the maze and sucrose rewards were delivered on either side of the maze. The animals’ behavior was first shaped to follow a figure-8 pattern, followed by two paradigms (**Figure 1F**). The first paradigm (“alternating”) required the mice to execute an alternating figure-8 pattern to receive a reward and the second paradigm (“left-only”) implemented a rule change in which the animal was only rewarded on the left side of the maze. Before initiating imaging in the alternating paradigm, the mice were fully trained to over 80% accuracy for three weeks. All mice were tested in both paradigms, starting with the alternating task, and followed directly by the change to left-only.

**Figure 1:**
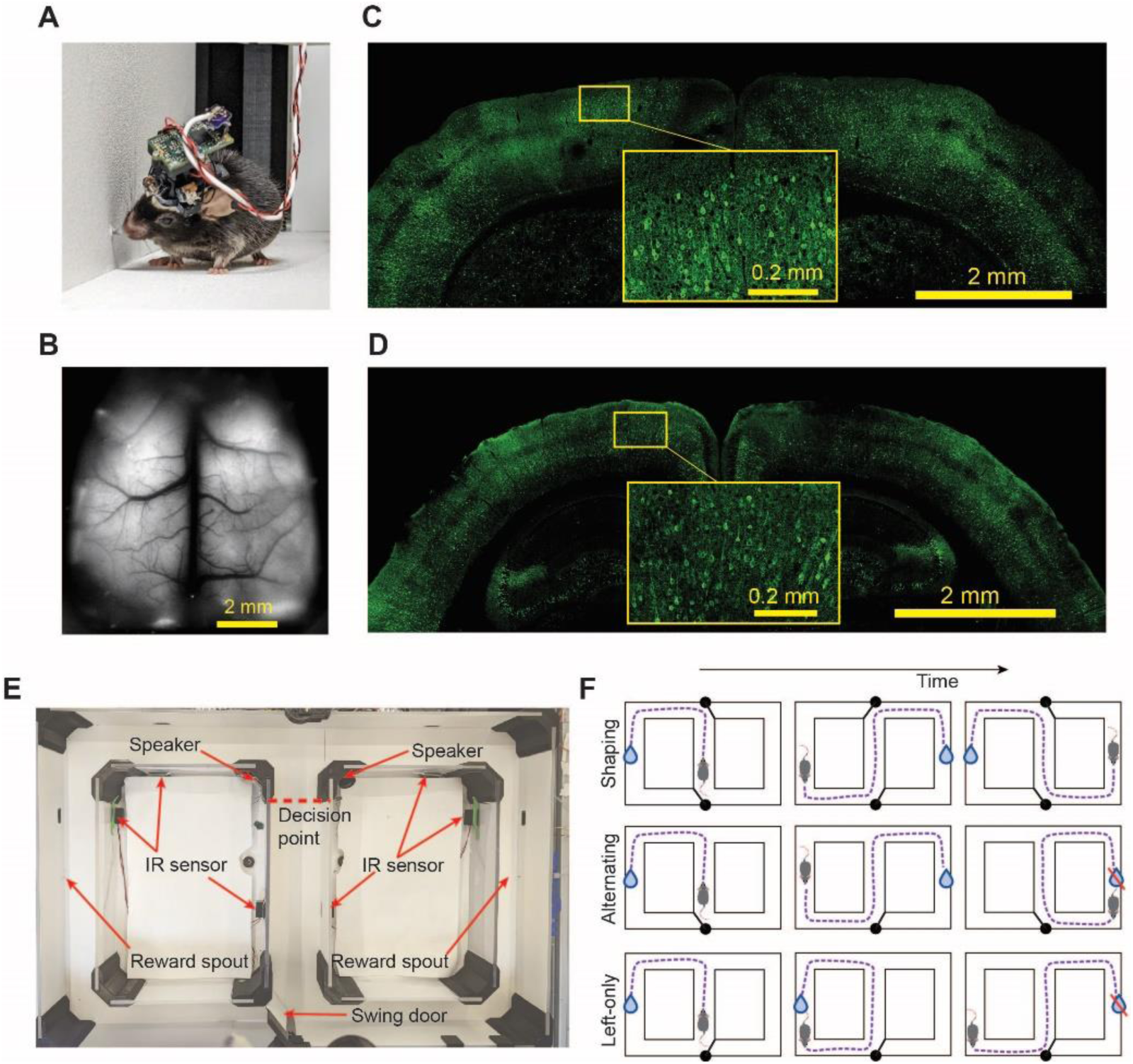
Widefield calcium imaging and the 8-maze apparatus. **A.** Mini-mScope attached to a mouse while in the 8-maze. **B.** Example brain imaging quality recorded from the mini-mScope. **C-D.** Coronal slices at 20x magnification stained for GFP from an anterior region (**C**) and posterior region (**D**). **E.** Overhead view of the 8-maze consisting of infrared break beam sensors that control the position of the motorized swing door and the delivery of a sucrose solution reward. **F.** Schematic of 8-maze behavioral paradigms. (Top) During the shaping phase, an automatic swing door was used at both the top and bottom of the maze to shape the animals’ movement in an alternating pattern. Each lap, doors swung in the opposite position to allow access to the opposite side of the maze. (Middle) The alternating phase utilized only 1 swing door so a free decision could be made at the top of the maze. Rewards were delivered only when an alternating pattern was met. (Bottom) The left-only paradigm rewarded only on the left side of the maze. however, the mouse had the ability to choose the right side and receive no reward.

### Behavior in the 8-maze

Mouse behavior during both the alternating and left-only paradigms was evaluated using a combination of manually scored and automatic pose estimation. Head position was used to determine instantaneous speed at each frame in the maze (**Figure 2A-B**). The average movement speed during correct alternating paradigm trials was significantly lower than during correct left-only paradigm trials (Alternating: 0.14 ± 0.01 m/s, Left-only: 0.18 ± 0.02 m/s, p = 0.03, t=2.98, df=5, Paired t-test; **Figure 2C**), consistent with additional time being allotted to decide on which maze arm to select. Across the 7 recording days of the alternating paradigm, the average preference for the left side of the maze was almost exactly half (50 ± 3.4%; **Figure 2D**), indicating no innate preference to one side of the maze over the other during this paradigm. After the left-only task was implemented, the mice developed a significant preference for the left side over the subsequent 8 recording days (left 73.2 ± 4.0%, p<0.0001, t=17.27, df =5, Paired t-test). Since the animals’ behavior was shaped to the alternating task prior to the recording trials, the decision accuracy was initially greater than 75% and remained stable for the next 6 recording days, with a slight decrease to 73% on day 7 (**Figure 2E**). Following the rule change to the left-only paradigm, decision accuracy was lower on the first day, followed by a steady increase in accuracy over the 8 testing days of the left-only paradigm. For the first 3 days of the left-only paradigm, accuracy was significantly lower than alternating (Day 1 p=0.0025, t=4.014, df=31; Day 2 p=0.0039, t=3.85, df=31; Day 3 p=0.0018, t=4.126, df=31, 2-way ANOVA with multiple comparisons, Bonferroni correction). However, the following 5 days revealed a steady increase in decision accuracy, comparable to the alternating task (**Figure 2E**). The alternating paradigm showed a significantly higher average decision accuracy than the left-only (**Figure 2F**, p=0.007, t=4.425, df=5, Paired t-test), due to the initial lower accuracy on left-only trials as the mice learned the rule change. Nevertheless, during both behavioral paradigms, the mice displayed an understanding of the task with total average accuracy above 70%.

**Figure 2:**
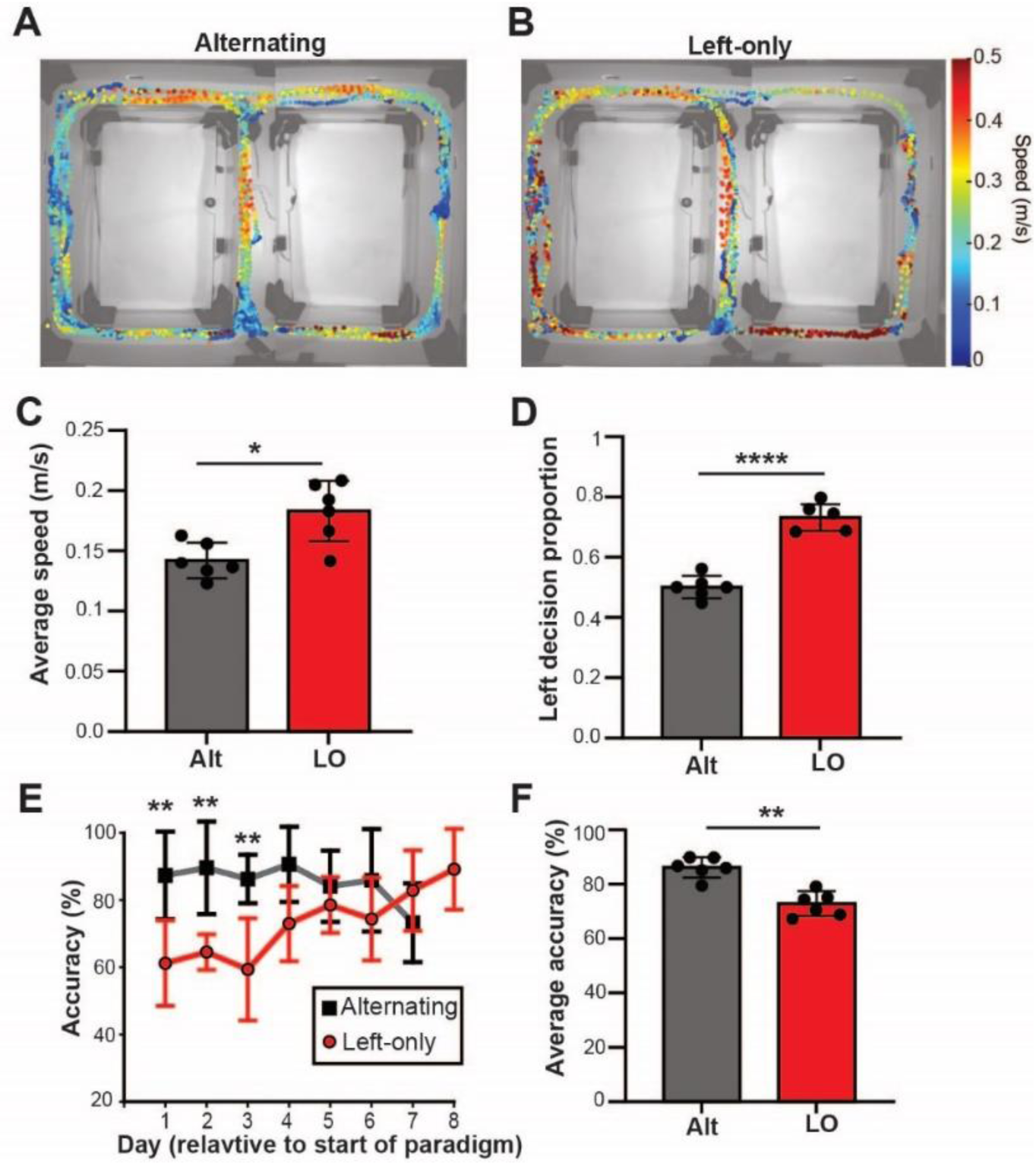
Behavior in the 8-maze. **A-B.** Head position during a representative 5-min alternating (**A**) and left-only (**B**) trial. Data points are color coded by instantaneous speed. **C.** Average movement speed ± SD per mouse for the two behavioral paradigms (Alt=alternating and LO=left-only). **D**. Average proportion of left decisions across all trials per mouse ± SD. **E.** Decision accuracy relative to testing day for both behavioral paradigms. **F.** Average accuracy across all trials for alternating and left-only (Mean ± SD).

### Cortical dynamics at key behavioral events

To assess patterns of Ca^2+^ fluorescence modulation of the cortical activity, we identified two salient behavioral events: making the correct decision and getting to the reward spout. Plots of the average z-scored ΔF/F for 7 CCF regions during a 6 s time window centered on either the moment the animal crossed the decision point or interacted with the spout during correct trials illustrate distinct, consistent activation patterns for both positions in the maze (**Figure 3A-B**). These CCF regions were selected because they represent diverse cortical areas as well as modulation in calcium fluorescence. At the time the mouse crosses the decision point (t = 0s), the activity of both motor regions (M1 and M2) decreases at -0.4 s. The somatosensory regions (SS_ll and SS_trunk) exhibit a local minimum average fluorescence at -0.4 s, followed by a general increase in activity. Similar decreases in activation at +0.13 s occur at the retrosplenial cortices (RSP_lat and RSP_dorsal) as well as earlier and later increases in fluorescence. Finally, a distinct increase in V1 activity precedes the decision point at -0.53 s.

**Figure 3:**
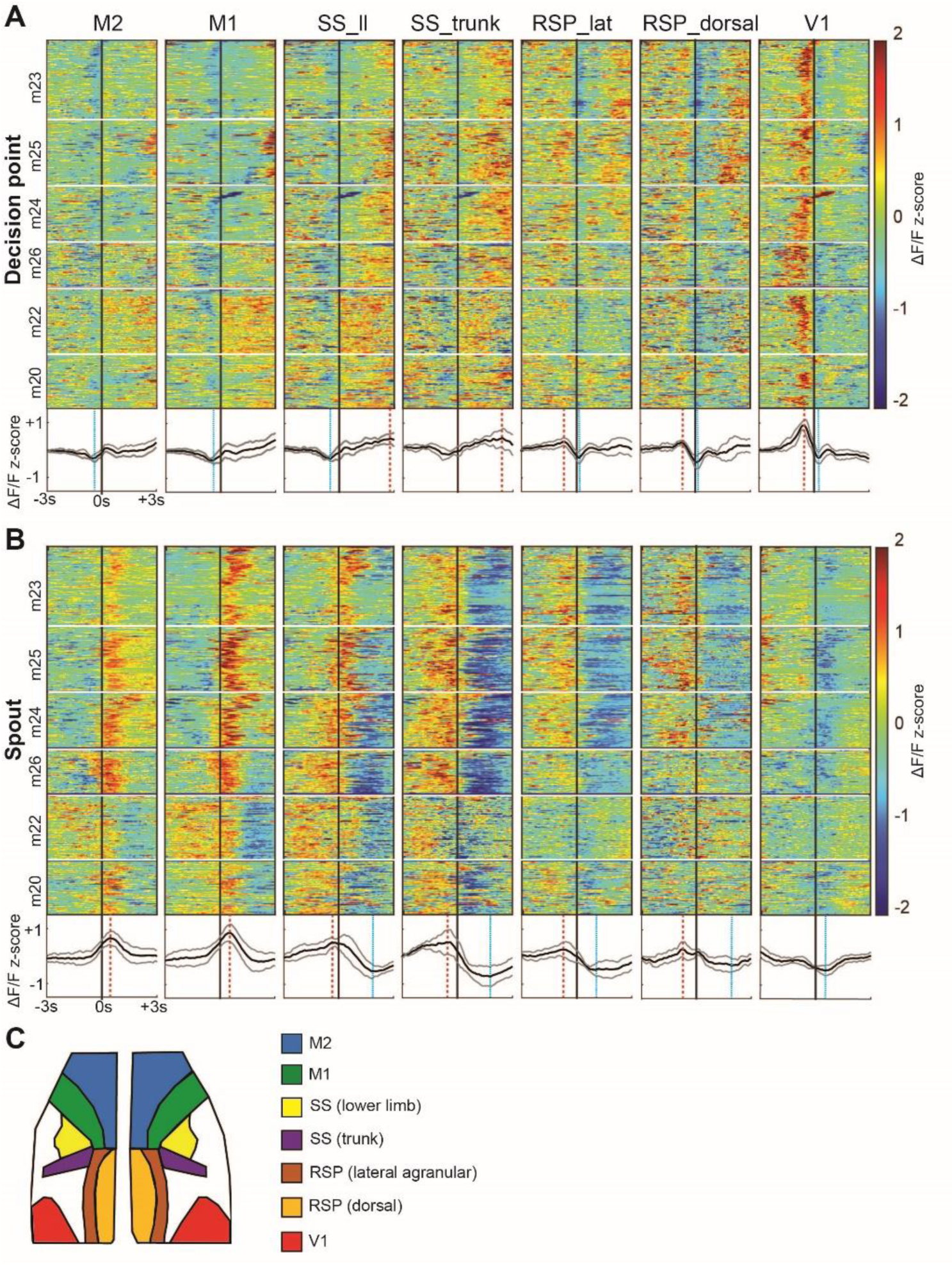
Cortical dynamics at key behavioral events. **A-B.** Plot of ΔF/F traces for all correct trials aligned to the time of crossing the decision point (**A**) and interacting with the spout (**B**). Individual mice are separated by horizontal white lines in the plots. Mean ± SD traces shown below each plot. Vertical red dashed lines indicate significant peaks compared to randomized data, dotted blue lines are significant troughs. Vertical solid black line indicates t=0 for the respective behavior. **C.** Atlas legend of the 7 CCF regions plotted in A-B.

When the fluorescence data is aligned to when the animals initially interacted with the spout (t=0 s), different activity patterns emerge. Immediately following interaction with the spout for correct decisions (**Figure 3B**), the motor regions show a prolonged increase in activity with peaks at +0.4 s. The somatosensory regions exhibit a biphasic activity pattern, peaking in activity prior to the spout at -0.4 s that reach a minimum after at +1.8 s. Retrosplenial cortical activity peaks at -0.7 s, followed by a decrease in activation. In contrast, V1 activation generally decreases around the spout interaction, with a minimum at +0.5 s. As described in the Methods and Materials, bootstrapping analysis demonstrates that all the average fluorescence peaks and troughs in **Figure 3A-B** exceeded the 95% confidence interval of the randomized data, so we conclude these are not chance events.

These ΔF/F signals were verified to reflect real cortical modulation and are not the result of confounding variables. First, to ensure the distinct cortical patterns following spout interactions were specific to the rewarding stimulus and not a confound of reduced movement speed, the average z-scored fluorescence at all stops outside of the spout region were compared with the peak fluorescence following correct spout interactions. Stops outside the spout regions were defined as any 1 sec segment where the animals average speed was <0.05m/s and was not within a defined region around the spout (**Supplemental figure 1A**). Statistical comparisons reveal that each CCF regions exhibited a ΔF/F signal that was unique to the reward spout, not reduced locomotion (**Supplemental figure 1B**, M2 p<0.0001, t=23.71, df=2358; M1 p<0.0001, t=28.33, df=2358; SS_ll p<0.0001, t=24.38, df=2358; SS_tr p<0.0001, t=17.81, df=2358; RSP_lat p<0.0001, t=13.71, df=2358; RSP_dor p<0.0001, t=9.187, df=2358; V1 p<0.0001, t=20.86, df=2358, Unpaired t-test, Bonferroni corrected alpha).

Next, to verify the patterns of activation in the central corridor were not an auditory confound from the tones (see Methods and Materials), we conducted alternating trials in the absence of tones and aligned the z-scored ΔF/F to the moment the animals crossed the center IR beam, where a tone would normally play. The average ΔF/F in each CCF region during alternating no-tone trials show similar patterns to those observed when the tones were present indicating these patterns were specific to the region of the maze, not simply a result of the tone (**Supplemental figure 1C**). Further, no-tone trials exhibit a clear activation of V1 at the IR beam followed by activation of somatosensory regions, matching the trials with a tone thus demonstrating the patterns of cortical activation measured are not an auditory confound. Finally, the average accuracy during tone and no-tone trials were nearly identical suggesting that the tones were not a meaningful factor in the animals’ decisions (Alt tone 0.86 ± 0.04; Alt no-tone 0.86 ± 0.09; p=0.96, t=0.05, df=5, paired t-test). With this, we do not make any further interpretations about how the tones effect decision behavior in the task, despite their presence in the 8-maze.

### Cortical activation states

While robust, repeatable ΔF/F modulation occurs at different CCF regions during behavioral events, the spatial activation patterns and their temporal sequences across the cortex are difficult to investigate in this manner. Therefore, a clustering method was used to classify each ΔF/F frame as one of a discrete set of cortical activation patterns to more accurately capture cortex-wide activations^28^ (see Methods and Materials and **Figure 4**). Using k-means clustering across all trials, an average of 12.2 (range 9-14) cortical activation patterns were identified across the 6 subjects, and 11 global clusters across all subjects (**Supplemental figure 2**). The spatial patterns of these clusters consist of localized regions of increased fluorescence, with a fraction of the remaining cortex having a relative decreased fluorescence. Across clusters, the regions of increased activation extend from the most anterior to posterior regions of the imaging field. Each ΔF/F frame was assigned to 1 of the 11 global clusters, which we refer to as cortical activation states, resulting in sequences of cortical activation states as a mouse navigates the maze. Aligning the cortical activation states to the moment the animals reached the decision point reveals patterns (**Figure 5**). For example, around the time of the decision point, activation of the V1/RSP state 8 (yellow) precedes activation of the M2/PPC state 11 (red). Many of these patterns of state activations are present across all mice. Qualitative differences between the two paradigms can be observed, with state 8 more prevalent prior to crossing the decision point during alternating than left-only trials (**Figure 5A-B**). Comparing the aligned cortical state data to the aligned ΔF/F time courses reveals similarities (see **Figure 3**), for example the strong activation of V1 preceding the decision point, verifying the k-means clustering extracts meaningful patterns from the ΔF/F data. When the state time courses are aligned to when the animals reach the spout, different sequences of states are evident, for example a prominent state M2 state 3 (blue), that also correlate with the ΔF/F time courses (**Supplemental figure 3**).

**Figure 4:**
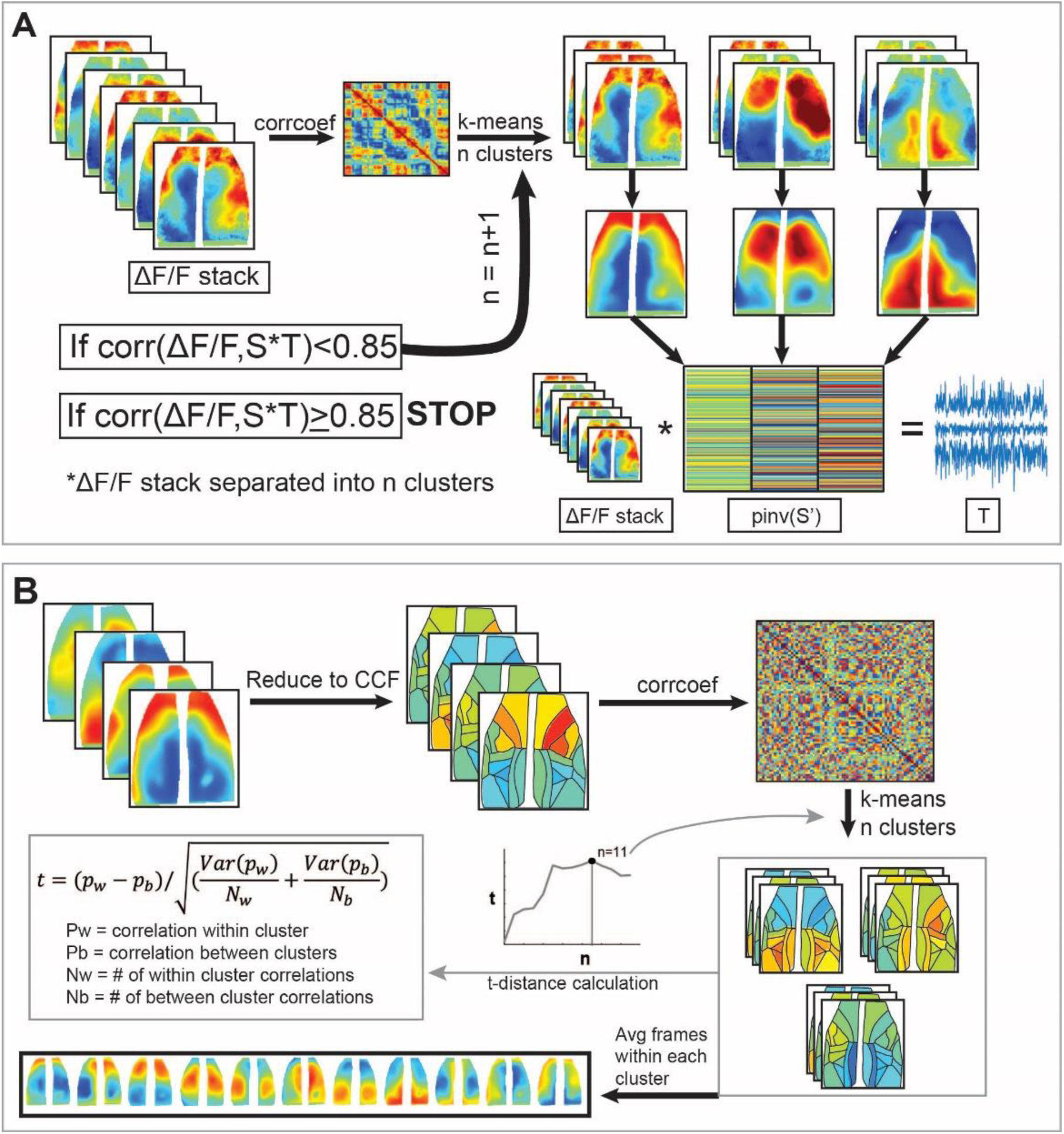
Identifying cortical activation patterns using K-means clustering. **A.** Trial-level clustering methods. ΔF/F stack represents all images collected in a 5-minute recording session (∼4500 frames). Correlation of each frame with all other frames is calculated, then clustered using k-means to determine within-trial cortical activation states. Optimal number of states determined with an 85% percent reconstruction threshold. Similar steps followed to cluster all trials for a single animal. **B.** Across-mouse clustering methods. Average cluster images are reduced to atlas regions, then correlation coefficients are calculated. Values are clustered using k-means and a t-distance algorithm was used to determine the optimal number of states. Final state images are determined by averaging all frames belonging to each of the 11 global states.

**Figure 5:**
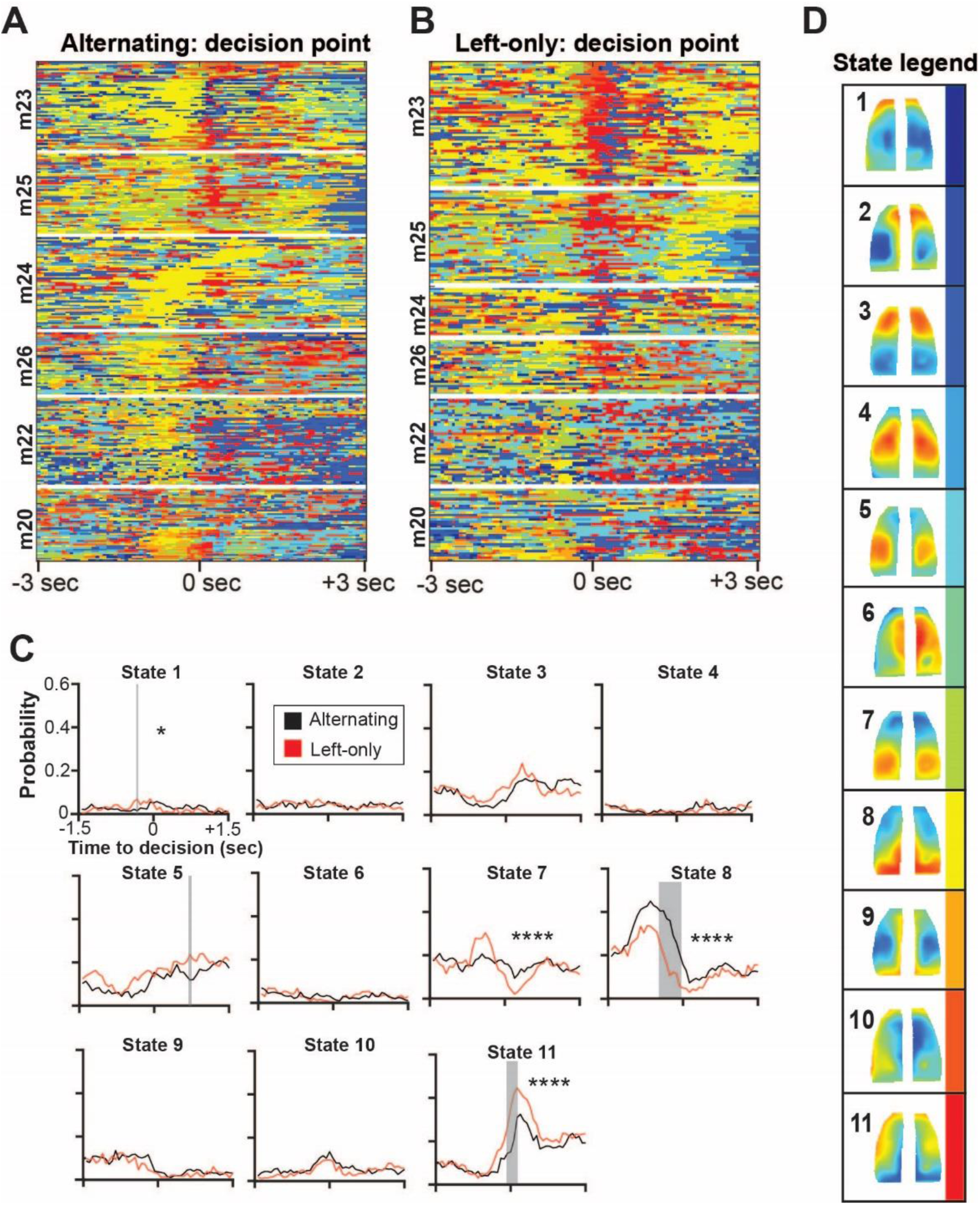
Cortical activation states aligned to the decision point. **A-B.** Cortical activation state time course ± 3 s with respect to the moment the mouse crossed the decision point during alternating trials (**A**) and left-only trials (**B**). Each correct lap is plotted by row across all 6 animals, with 470 total correct laps. For state time course aligned to spout interaction, see **Supplemental Figure 3**. **C.** Peri-event probability curves for each state with respect to the decision point. The probability of each state activating was calculated and compared between the alternating (black) and left-only (red) paradigms. Post-hoc time points where the two paradigms are significantly different are highlighted in gray. Main effect significance is indicated with (*).

As the two paradigms require different numbers of left and right decisions, we tested for differences in the state time courses for left and right-hand spout interactions during alternating trials alone (**Supplemental figure 1D-E**). Comparison of the average probability curves for each state reveals a main effect difference between the left and right spouts for 3 of the 11 states (**Supplemental figure 1F-G**, 2-way ANOVA with multiple comparisons, Bonferroni correction, State 1: p=0.0003, F(89,445) =1.703; State 2: p=0.011, F(89,445)=1.432; State 11: p=0.046, F(89, 445) = 1.302). However, post-hoc testing did not identify any significant time points to describe these main effect differences. Therefore, we interpret any state activation differences between the sides of the maze to be trivial, and any major changes in state probabilities are related to the paradigm.

To test for differences in state activations between the two behavioral paradigms, the peri-event activation probability curves for a 3 s window centered on the moment the mouse reached the decision point were calculated for each state and compared across behavioral paradigms (**Figure 5C**, see Methods and Materials for detailed description of statistical comparisons and exclusion criteria). State 8 has a significantly higher activation probability during the alternating paradigm (F(45,225)=2.95, p<0.0001), suggesting this task requires greater utilization of cortical regions involved in visual processing and spatial navigation. Alternatively, state 11, a combined M2 and PPC state, has significantly increased probability during left-only trials near the decision point (F(45,225)=2.38, p<0.0001), indicating greater involvement of motor planning in M2 in concert with the PPC as the mice traverse the maze to obtain the reward. We also compared state probability curves during the spout interaction (**Supplemental figure 3C**), finding the only significant difference is an increased activation probability of the RSP/V1 state 8 during the left-only trials prior to reaching the spout (F(45,225) = 2.552, p<0.0001). Since the spout interaction occurs following the decision, the decreased probability of state 8 during the alternating paradigm may indicate that more cortical regions used in working memory are engaged even after the decision during this paradigm.

### State activation frequency by behavioral region

To understand the relationship between cortical state and maze location, we determined the probability of each state activation when the animal was within the 7 maze regions (**Figure 6A-B**). These maze regions, which are symmetrical on the two sides, can be used to segment the task into 4 behaviors: decision-making in the central corridor, approaching the reward spouts, receiving the reward, and returning to the central corridor. By comparing the actual probability of state activation by maze region with shuffled data (alpha <0.0001, see Methods and Materials), we identified 25 significant state probabilities across the 7 maze regions (**Figure 6C-D**). Consistent with the state activation plots (**Figure 5**), state 8 has a high probability of activation before the decision point in the central corridor and state 11 has a high probability after the decision point in regions 2 and 5 along the path to the reward. For both paradigms, the probability of the M1/M2 state 3 increases in the two spout regions (regions 3 and 6). As this activation occurs in the right spout region where no reward was delivered during left-only trials, this suggests that state 3 encodes anticipatory and/or error information.

**Figure 6:**
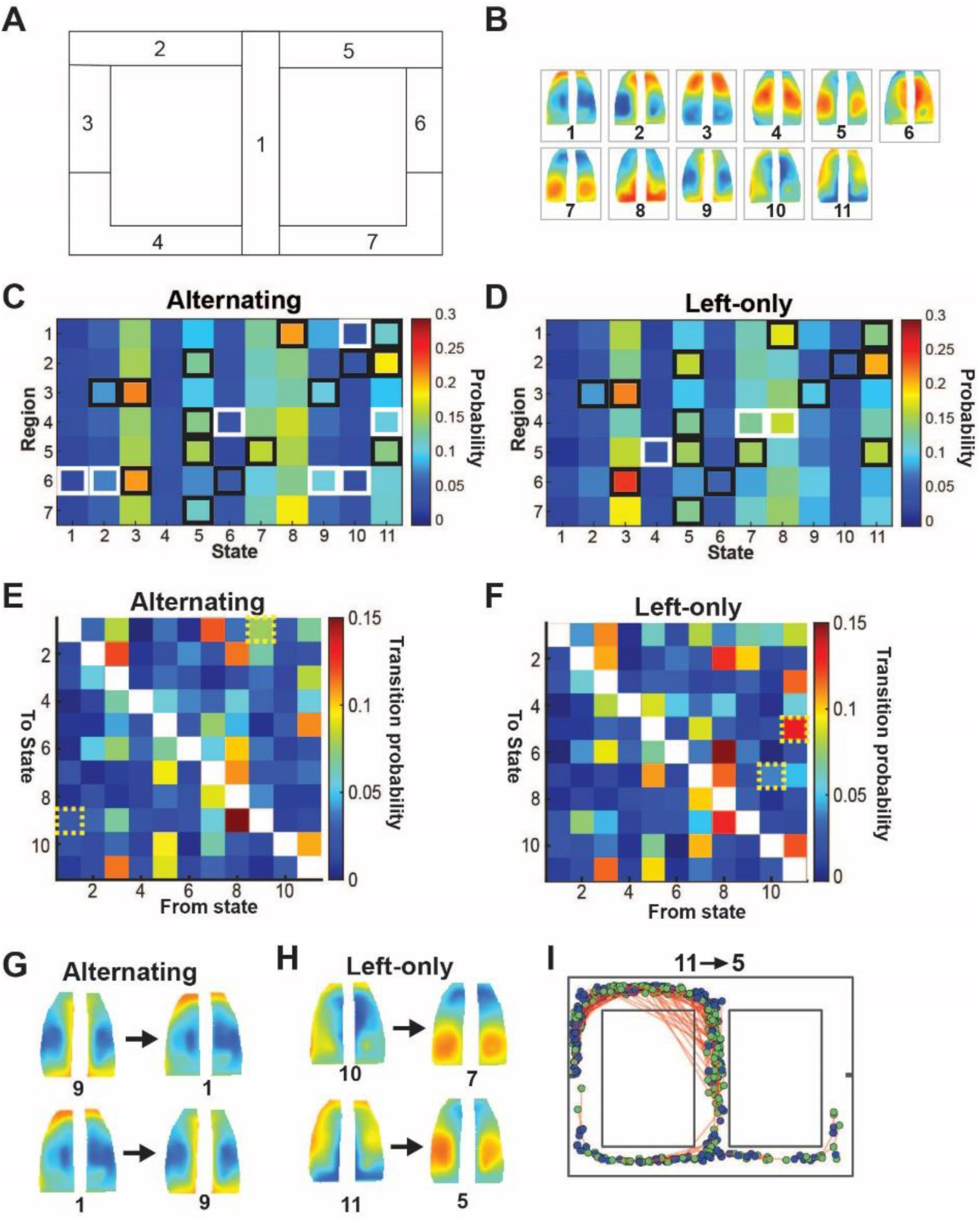
State probability by region and state transitions. **A.** Legend for the 7 maze regions. **B**. Legend for the 11 cortical activation states. **C-D.** State activation probability for each maze region during the alternating paradigm (**C**) and left-only paradigm (**D**). Black boxes indicate probabilities significantly greater than chance (>mean+5 SDs from randomized) that are present during both paradigms. White boxes indicate significant probabilities unique for the respective paradigm. **E-F**. Probabilities of each cortical activation state transitioning to every other state during the alternating paradigm (**E**) and left-only paradigm (**F**). Significant transition pairs shown in yellow dotted boxes. **G-H**. Average state images corresponding to the significant transitions during the alternating paradigm (**G**) and left-only paradigm (**H**). **I.** Mouse locations for the 11,5 transition during the left-only paradigm. Green dots indicate location at state 11 onset and blue dots indicate position at state 5 offset. Onset and offset pairs are linked with red line.

### Cortical two state transitions

In addition to the cortical activation state probabilities as a function of maze position, an analysis of the two state transitions assessed how information is passed in the cortex during decision-making behavior. Using the state data of a 6 second time window centered on crossing the decision point, the probability of state x transitioning to state y was calculated, resulting in a 11x11 matrix for both the alternating and left-only paradigms (**Figure 6E-F**). Self-self state transitions (matrix diagonal) were omitted as they do not describe state transitions. The probability of each state transition was compared with bootstrapped, shuffled data to determine significance (alpha < 0.05, see Methods and Materials). During alternating trials, significant transitions include the RSP/V1 state 9 to frontal state 1 and the reverse transition (**Figure 6G**), suggesting feedforward and feedback communications between frontal and visual/RSP regions. During the left-only task, significant transitions involve frontal states 10 and 11 to posterior states 7 and 5, respectively (**Figure 6H**). These left-only transitions suggest a feedback mechanism by which frontal and lateral regions influence PPC and V1 processing of sensory information. More generally, these two state transitions demonstrate the occurrence of directional flows of cortical information, with posterior to anterior (state 9 to 1) and anterior or anterior/lateral to posterior (states 1 to 9, 11 to 5, and 10 to 7).

To understand where these two state transitions occur in the maze, we plotted the onset and offset locations of these four significant transitions. The 9,1 and 1,9 transitions occur in similar locations in the maze during the alternating paradigm with onset and offset locations throughout the central corridor and following the decision point, predominantly on the left side of the maze (**Supplemental figure 4**). The 10,7 transition shows similar distribution both before and after the decision point during the left-only paradigm. The 11,5 transition shows a higher frequency of expression with state 11 onsets concentrated prior to the decision point and state 5 offsets after rounding the corner (**Figure 6I**). Therefore, during the left-only task, anterior to posterior cortical communication between M2/PPC and visual regions are engaged while making the decision.

### Cortical activation state motifs

To provide further insights into the flow of activation across the cortex as the mouse navigates the maze, we characterized temporal sequences of cortical activation states. Referred to as cortical state motifs, we identified 3-state sequences that were both statistically different from randomized sequences and exceeded a minimum probability of occurrence (see Methods and Materials). This resulted in 23 significant state motifs during alternating trials, and 30 during left-only trials, with 13 common motifs across both paradigms. Among these significant motifs,14 are characterized by a primarily anterior propagating sequence of activations and 13 by a posterior propagating sequence (**Figure 7A**). In addition to commonly expressed motifs, both paradigms display unique anterior and posterior propagating motifs.

**Figure 7:**
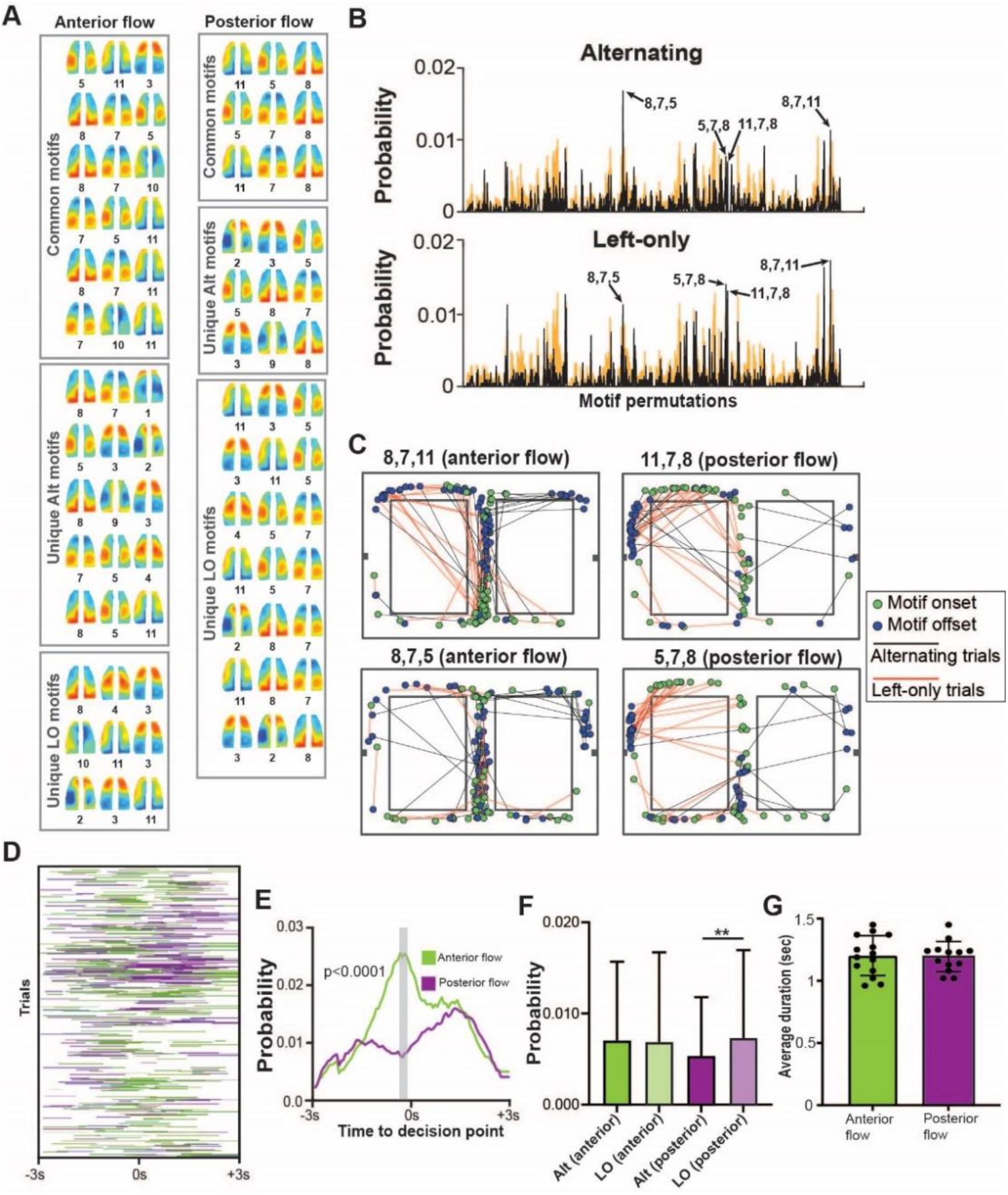
Cortical state motifs. **A.** All significant 3-state motifs separated into anterior and posterior propagation patterns. Unique alternating (Alt) and left-only (LO) motifs were found to be significantly more likely than chance for one paradigm but not the other. Common motifs were significant from chance during both paradigms. **B.** Probabilities of all state motif permutations. Real probabilities shown as black line. Bootstrapped probabilities +5SD shown in orange. Specific motifs of interest are highlighted with arrows. **C**. Representative plots of mouse locations during 2 common forward propagating motifs and their respective reverse motifs. Onset locations shown as green points and offset locations shown as blue points. Behavioral paradigm is signified by the line color connecting the onset and offset points (black=alternating; red=left-only). **D.** Plot of anterior propagating and posterior propagating motif activations during a 6-second time window centered on the decision point for each correct trial (Green=anterior flowing motifs, Purple=posterior flowing motifs). **E.** Average motif activation probability curve over the same time window from (**D**). Average anterior propagating motif curve shown as green line, average posterior propagating motif curve shown as a purple line. Significant post-hoc points highlighted in gray area. **F**.

Rather than randomly distributed probabilities across all possible motif permutations, there are peaks in probability for specific motifs (**Figure 7B**). Both specific sequences and propagation characteristics highlight the differences in the two paradigms. Comparing the probabilities of the most prevalent motifs for both paradigms, 8,7,11 represents a complete posterior to anterior propagation along the cortex during the left-only task, while the 8,7,5 motif a partial propagation for the alternating task (**Figure 7B**). Anterior motifs are active for 16.5% of the 6-second time window centered on the decision point across all trials, and posterior motifs are active for 11.6% of the time. Across all 6 subjects, anterior motifs are expressed in 55.2 ± 9.3% of correct trials, and posterior motifs in 24.8 ± 14.4%. Thus, around the decision, these directional flows of cortical activity are both commonly expressed and found in all mice.

Plotting the mouse head position at the times of onset and offset for some of the more prevalent motifs provides qualitative evidence that anterior and posterior propagating motifs are utilized at different positions along the maze. For example, anterior propagating motifs for the V1/RSP state 8 (8,7,11, or 8,7, 5) have onset locations at the bottom and top of the central corridor with offset locations either at the top of the central corridor or as the mouse rounds the corner to either reward spout (**Figure 7C**). The respective reverse motifs (11,7,8 and 5,7,8) have a different spatial distribution, with onsets occurring before and after the decision point, and offsets clustered just prior to the spout.

We then plotted each activation of the anterior propagating and posterior propagating motifs during the 6 s time window centered to the decision point (**Figure 7D**) and calculated the average peri-event probability for all motifs for either direction with respect to the decision point. Anterior propagating motifs have peak probability near the moment the animals cross the decision point, and posterior propagating motifs have highest probabilities around 1.5 seconds prior to and following the decision point (**Figure 7E**). This result suggests that sequential feedforward and feedback mechanisms are engaged as the mice navigate and make decisions along the maze. Comparison of these average curves reveals significantly different expression probabilities among the two flow directions (p<0.0001, F(89,2225)=2.623, 2-way ANOVA with multiple comparisons, Bonferroni corrected). By comparing the average probability of the 14 anterior motifs and 13 posterior motifs during the two paradigms, there was no significant difference between paradigms for anterior motifs (p=0.86, t=0.17, df=83, paired t-test), but a significant increase in posterior flowing motifs during the left-only task (p=0.028, t=2.25, df=77, paired t-test, **Figure 7F**). Therefore, anterior propagation of activity is necessary for decision generation and maze navigation during both paradigms, while posterior propagation is utilized more often by the left-only paradigm further supporting that unique cortical feedback patterns arise when the load of working memory is reduced.

Next, we determined activation duration for all anterior and posterior motifs during a 6 s time window relative to the decision point for each correct lap. The average durations of the anterior and posterior propagating motifs are nearly identical (1.20 ± 0.16 sec, 1.19 ± 0.12 sec respectively, **Figure 7G**, p=0.88, t=0.1496, df=25, Unpaired t-test) indicating the differences in utilization of forward and posterior motifs cannot be simply explained by duration of activation. Instead, these motifs suggest a cortex wide temporal parcellation scheme that directs task-dependent information in two directions, with the anterior propagation primarily feedforward information and the posterior propagation primarily feedback.

## Discussion

We used widefield Ca^2+^ imaging to record neural activity during an 8-maze decision task to investigate differences in cortical involvement during two variants of the task. Since activation of specific cortical regions during goal-directed navigation is dependent on task context^32^, and cognitively-demanding tasks utilize more spatially diverse cortical regions^13^, we hypothesized that the cortical circuits engaged will depend on the degree to which working memory is required. We implemented a k-means clustering algorithm to characterize spatiotemporal patterns of cortical activation, referred to as cortical activation states^27,28^, and calculated probabilities of these states as well as sequential combinations of multiple states. These findings support the theory of a bidirectional perception-action cycle by highlighting how the cerebral cortex dynamically modifies how information is passed among regions to most effectively execute decision-making behavior.

### Spatiotemporal patterns of cortical state activations

The cortical state probability curves demonstrated significant differences in state activation patterns during the decision phase of either behavioral paradigm. State 8, characterized by activations in V1 and RSP, has a significantly higher activation probability during alternating trials than during left-only trials. In contrast, a M2/PPC state 11 has an increased probability during the left-only paradigm. These changes in state utilizations across behavioral paradigms were observed both with the peri-decision probability curves (**Figure 5C**) as well as the state probabilities in maze regions (**Figure 6C-D**). The peak probability of V1/RSP (state 8) activation prior to the decision point in the central corridor during the alternating paradigm likely reflects a difference in how visual information is utilized during memory recall under this paradigm. During the alternating task, the required knowledge of the previous decision may increase engagement of widespread cortical regions that initiate an anterior flow of activation beginning at the most posterior cortical regions. Given the roles of RSP in spatial navigation and spatial memory^33,34^, a task involving memory of recent spatial location will also likely require greater use of this cortical region.

The reduced probability of the V1/RSP activation (state 8) and increase of M2/PPC (state 11) during the left-only paradigm suggests an alteration in cortex-wide involvement during the decision-making process. To explain this difference, at least partially, this result is consistent with the role of frontal cortical regions in establishing reward contingencies^35–39^. As an important neural hub for integrating decision and reward information, M2 contributes to the determination of the best decision given the possible outcomes^40,41^. The increase in state 11 suggests that the action-reward contingency is more defined during the simplified left-only paradigm. Furthermore, the average movement speed was higher during the left-only compared to alternating trials (**Figure 2C**), suggesting a reduced cognitive load on the animals when generating a decision and a more efficient transformation of perception to the correct decision. Given the PPCs involvement in the transformation of sensory inputs to motor regions to guide movement and make decisions^42^, the increased use of state 11 during the left-only paradigm may reflect the initiation of a simplified flow of activation that starts at slightly more anterior regions, rather than at V1. Perhaps the perception-action circuit during the left-only paradigm relies less on the processing of visual and spatial information and shifts the anterior flow of activation forward.

### Cortical two state transitions and motif flow patterns

The dorsal stream is a long-standing tenet that visual information is processed sequentially in higher-order brain regions as it is passed in the anterior direction^2,43–45^. More generally, this posterior-anterior flow of information is implicated in mediating the transformation of sensorimotor information during sensory-guided tasks^2^. Wide-field calcium imaging has shown that both the spatial and temporal pattern of posterior to anterior activation is refined during learning of an auditory task^46^. More recently, anterior to posterior propagations of information have been hypothesized to be involved with various cognitive functions^1,3,6,47,48^. From these perspectives, we sought to identify differences in two state transitions between the two paradigms. During the alternating paradigm, transitions between RSP state 9 and M2 state 1 are significantly more likely than chance, consistent with task demands and involvement of RSP in spatial memory and spatial navigation^33,34^. The combination of these mirrored transitions reflects the engagement of the reciprocal circuit between M2 and RSP^49^ that might be necessary during the recall of previous decisions. During left-only trials, transitions from M2 states 10 and 11 to visual and somatosensory states 7 and 5, respectively (**Figure 6E-H**), were significantly greater than chance, suggesting increased anterior to posterior interaction during this paradigm. As described above, V1/RSP state 8 shows more pronounced activation during the alternating paradigm. Interestingly, this increase in transitions from frontal to posterior states including V1 and PPC occurs in left-only, not alternating trials. The seemly incongruence of these two results can be explained by the direction in which the activation flows. The higher frequency of these anterior-posterior transitions (states 11 to 5, and 10 to 7) suggests the left-only paradigm involves more internal feedback to guide subsequent behavior. Because the animals displayed a significant increase in average movement speed during the left-only paradigm, this increased utilization of feedback may contribute to a more efficient generation of the decisive action. These differences in two state transitions between paradigms highlight an additional mechanism on how cortical dynamics can change after implementation of new rule to match altered task demands and/or behavioral strategies.

The state motifs analysis further characterized these directional flows of activation, finding that flows in the anterior and posterior directions are common. Therefore, while we observed the generally accepted flow of information from sensory cortices to PPC and to frontal cortices, the results demonstrate that the previously hypothesized posterior flows also occur as the mice navigate the maze. There are differences in both the probability of common states as well as the state motifs between the two paradigms. For example, comparison of the most prevalent motifs between the two tasks suggests the complete posterior to anterior transformation of information described by the 8,7,11 motif is more common during the left-only task, while the partial transformation shown in the 8,7,5 motif is more common for the alternating task (**Figure 7B**). Previous studies found that once a task is learned, the sequence of cortical activity to execute movement becomes temporally compressed, indicating a more efficient utilization of task information^19^. Perhaps the decrease in cognitive resources required during the left-only task allows for fewer steps in the progression of information anteriorly reflecting a more efficient sensorimotor transformation in a single motif. With more steps required for the alternating paradigm, as it relies on working memory.

These spatiotemporal sequences of activations not only provide new evidence that information flows through the cortex in both directions, but also that these anterior and posterior flows occur at distinct times during the perception-action cycle. Anterior propagating motifs are most likely to occur immediately before crossing the decision point, suggesting engagement of the dorsal stream pathway as the animal processes sensory information in the central corridor. Conversely, the peak probability of posterior propagating motifs occur after the decision was made, similar to the two state transitions. Perhaps this peak in posterior propagating motifs following the decision, reflects an internal feedback mechanism by which higher-order regions of the frontal cortex project to posterior sensory regions to update an internal model on task performance and prepare for or guide future actions, for example the upcoming reward. Most of the posterior motifs involve M2, consistent with both its intracortical connectivity and numerous roles in cognitive and executive processes. The latter include engagement in perceptual tasks, decision-making and choice, executing and learning complex movements, spatial navigation, and feedback to predict optical flow (for review see^4^). These roles have led to the proposal that M2 provides top-down information about these processed to cortical and subcortical regions^4,6^, consistent with the posterior flows observed.

Interestingly, the average duration for the anterior and posterior flow motifs were almost identical at 1.2 s, indicating a uniform and relatively slow bidirectional movement of information. Comparing flow direction during the two paradigms reveals that an anterior flow is necessary during both paradigms, further supporting the classic interpretation of sensory information to action via the dorsal stream. However, posterior flow motifs are significantly more likely during the left-only paradigm providing evidence that the left-only paradigm uses a different cortical processing scheme that relies more on internal feedback for future action.

### Conclusions

In conclusion, we performed widefield Ca^2+^ imaging in freely moving mice navigating two versions of an 8-maze and uncovered well-defined patterns of cortical activation at key behavioral aspects of the task. Differences in the probabilities of cortical activation states, two state transitions, and sequences of states distinguish the alternating and left-only reward paradigms. The transformation of visual and spatial information into decision-making involves the integration of more spatially diverse cortical regions during a task requiring working memory of the previous decision. Different probabilities and sequences of cortical state activation also characterize the paradigms. The alternating task utilizes patterns of activation flow from posterior to anterior including extensive involvement of the visual areas, RSP, PPC, and M2. The left-only paradigm involves a higher frequency of posterior flows of activation from M2 to PPC, suggesting a simplified sensorimotor circuit. While the posterior to anterior flows provides widefield imaging support for the widely accepted dorsal steam hypothesis, the demonstration of equally prominent reverse flows emphasizes the importance of directional information exchange across the cortex during the decision-action cycle.

## Methods and Materials

All experiments were approved by the Institutional Animal Care and Use Committee (IACUC) at the University of Minnesota.

### 8-Maze and behavior task

Based on these previous modified T-maze designs^50–56^, we constructed an automated 8-maze that was able to detect mouse position and deliver a drop of 5% sucrose solution on either side of the maze after a desired rule was followed. The 8-maze was an 80 x 54 cm rectangular arena constructed using clear acrylic and custom 3D-printed connection pieces (Creality CR-10s, HATCHBOX 1.75 mm PLA) (see **Figure 1E**). The walls of the arena were 12 cm tall, and corridors were 8 cm wide. The internal walls and floor of the maze were covered with a white, textured vinyl sticker to create traction for the mouse and promote higher visual contrast. Intersections at the top and bottom of the center corridor had motorized swing doors to shape the mouse’s path in an alternating path during behavior training sessions. Using an Arduino script, servo motors (Aideepen, MG996R) and infrared break beam sensors (Adafruit Industries, #2168) controlled the position of 3D printed swing doors relative to the mouse’s position in the maze. To reduce potential IR penetrating light artifacts, break beam sensors were covered with a vinyl sticker that had a pinhole to allow just enough light through for the sensor to function.

Two variants of the 8-maze task were implemented to target different aspects of cognitive processing and examine for different patterns of cortical activation. The first paradigm required the mice to execute an alternating figure-8 pattern to receive a reward (**Figure 1F, middle**). A solenoid valve was used to deliver a 27 µL reward of 5% sucrose through a rubber tipped needle spout on either side of the maze when the correct decision was made (The Lee Company, #LHDA0533315H). The second paradigm implemented a rule change where the animal was only rewarded on the left side of the maze (**Figure 1F, bottom**). During this left-only paradigm, the right side of the maze remained open to exploration but did not deliver a reward. The distance the animals were required to travel was equal for both paradigms, and each side of the maze was identical. All mice were tested in both paradigms, starting with alternating, and followed directly by the change to left-only. This way, any mouse-specific changes in behavior could be isolated and the differences in cortical activity associated with the rule change could be determined.

Speakers were mounted on the maze walls on either side of the decision point (**see Figure 1E**) and played one of 3 innocuous tones at specific times during testing. During the alternating paradigm, a brief 3520 Hz tone played on the right speaker when the mouse was in the central corridor to signify a right-hand turn is necessary for a reward and a 1047 Hz tone was played on the left speaker when a left-hand turn was necessary for reward. 15 cm after the decision point on either side of the maze a 2637 Hz reinforcement tone was played only when the correct decision was made. Tones remained consistent for the left-only task, with the only difference being that only the 1047Hz was played in the central corridor to signal only for left-hand turns. Two behavior cameras were mounted 1.5 m above the maze (30fps, FLIR blackfly). Prior to each trial, the maze was thoroughly cleaned with 70% ethanol to eliminate inter-trial olfactory cues. The entire maze was enclosed in an arena with black curtains hanging on each of the walls to reduce the amount of light in the maze. Only ambient room light from underneath the maze was present during testing trials.

Mouse training in the 8-maze lasted for 3 weeks, broken into separate 1-week phases. The first phase simply involved handling the mouse for a 5-minute session per day. The second phase consisted of placing the mouse in the 8-maze with a weighted dummy head-mounted microscope and providing a sucrose reward whenever the mouse reached the spout. During the third phase, a functional mini-mScope was attached, including the data and power wires. Mice were then taught to follow a figure-8 alternating pattern to receive a sucrose reward using both automatic swinging doors (5-minute trials per day). To verify the behavior was learned, the door at the top of the maze was removed to give the mouse free choice, and decision accuracy was measured. After shaping was completed and the mice achieved a decision accuracy ≥75%, cortical Ca^2+^ imaging was initiated during the alternating task for 7 days, followed by the left-only task for 7-8 days. Only a single recording session per day per mouse was run to ensure the animals were consistently water deprived and not satiated by the water delivered from the reward spouts.

### Behavior analysis

Mouse behavior in the 8-maze task was analyzed using a combination of manual scoring and automatic pose estimation software (DeepLabCut)^57^. Manual scoring consisted of finding the exact frame where the animal’s nose crossed the decision threshold defined in **Figure 1E**, and the frame when the nose reached the reward spout during correct trials. If the mouse did not stop at the spout, this frame was defined as the point where the nose is closest to the spout for the particular lap. Using DeepLabCut, the position of the mouse’s nose, head, body, and base of the tail were tracked and quantified. For speed and position analysis, the head position was used as it had the most accurate measure of tracking.

### Animals, cranial windows, and viral vector injections

In this study, 6 C57BL/6 mice (2 male, 4 female, average age of 225 days at first day of testing) were used. Animals were housed in a 12-hour reverse light/dark cycle and all behavioral testing was done during the dark phase. Mice were water restricted to 1 mL per day and were maintained at a weight at least 80% of their pre-restriction weight. The mice were implanted with a version of our previously described cranial polymer window^16,17,27,28^. Following a 1 week recovery from implantation, the mice were then retro-orbitally injected^58^ with 4.4×10^11^ genocopies of a GCaMP7f viral vector in 100µl of sterile saline (AAV-PHP.eb-Syn-jGCaMP7f-WPRE; Addgene #104488). The viral vector incubated for at least 3 weeks prior to Ca^2+^ imaging recordings.

### Immunohistochemistry

To harvest the brains for tissue processing, the animals were deeply anesthetized using 5% isoflurane and transcardially perfused first with ice-cold phosphate buffered saline (PBS) followed by 4% paraformaldehyde (PFA). Extracted brains were fixed in PFA for 2 days before sectioning at 50 µm thickness using a vibratome. Brain slices were then gently shaken in a blocking solution (10% Donkey serum in 0.1M TBS) for 1 hour. The slices were next incubated in a 1:1000 dilution of the primary antibody for GFP staining (rabbit anti-GFP, Invitrogen, REF# A6455) overnight at 4°C followed by three 10-minute washes in tris buffered saline (TBS). The slices were then incubated in 1:1000 dilution of the secondary antibody overnight at 4°C (Donkey anti-rabbit Alexa Fluor Plus 488, Invitrogen, REF# A32790). Using a confocal microscope (Leica Stellaris 8), images of the sections were taken at 20X magnification with a 488 nm laser. The images were then tiled together to get a single high-resolution image for each section (see **Figure 1C-D**).

### Widefield imaging with the mini-mScope

A miniaturized head-mounted microscope (mini-mScope) was used to record Ca^2+^ activity during both the alternating and left-only trials in the maze (n = 34 alternating, n = 29 left-only, from 6 subjects). The mini-mScope body was adapted from a previous design^27,28^ and 3D printed from polymethyl methacrylate (PMMA) resin using a stereolithography printer (FormLabs). To ensure that no light could pass through the PMMA, an additional coating of black paint was applied to the exterior (DYKEM BRITE-MARK). Underneath the mini-mScope body, 3 magnets were inserted for attachment to the magnets in the cortical window implant. The body housed 4 blue LEDs (470 nm, Digikey) radially distributed around a central shaft and pointed down to the dorsal surface of the cortex. To eliminate wavelengths of light outside of the range that excites GCaMP7f, bandpass excitation filters were added for each LED (450-490 nm, Chroma). A miniaturized imaging sensor, the miniFAST was adopted from previous work^59^ to be used for Ca^2+^ imaging. The housing for the CMOS sensor was machined from polyoxymethylene (Delrin). The focal plane was adjusted by sliding the CMOS housing along the mini-mScope shaft until a clear brain image was displayed, and a screw was used to anchor the CMOS in place. The emitted light from GCaMP7f was directed to a collecting lens in the central shaft of the mini-mScope body and was passed through a bandpass emission filter (500-550 nm, Chroma) before reaching the CMOS sensor.

### Widefield Ca^2+^ recordings

The CMOS sensor was set to record at 30 frames per second (fps) with the gain set to 25 for a total trial duration of 420 seconds. To achieve stable LED power and mitigate heating issues, the LEDs were flashed at 15 Hz for 17 ms using TTL pulses generated from a microcontroller (Teensy 3.5). The data acquisition unit (DAQ) for the miniFAST CMOS sensor synchronized the behavior cameras with the CMOS sensor also using TTL pulses. Prior to each recording trial, the PET window on the animal’s implant was gently cleaned with saline to remove any dust or debris from the imaging field. During the first 120 s of each trial, no Ca^2+^ data was collected to allow the LED light power to stabilize. Following this stabilization period, the mini-mScope was attached to the mouse and the animal was placed in the center lane of the 8-maze facing the decision intersection. During the recordings, a manually controlled motorized commutator was used to relieve torsion in the coaxial and LED power cables^60^.

### Ca^2+^ imaging data pre-processing

All data processing was completed using custom MATLAB scripts (MATLAB 2022b). The 30 fps recordings were first down sampled to 15 fps to remove all frames in which the LED was off. The video was then rotated 180 degrees to have the rostral cortex at the top of the image. Each frame was converted to grayscale and resized using an 80% bilinear compression. A mask was drawn to exclude pixels in the background and central sinus. The ΔF/F calculation utilized a previously described method that normalizes the data to changes in global illumination fluctuations to account for differences in LED power during each trial^61^. Each pixel of the image stack was divided by its mean over the entire recording. From this, the mean fluorescence of each frame divided by the mean of the entire stack was subtracted yielding a normalized image stack. The data were spatially filtered by averaging a 14 pixel-radius and temporally filtered with a 0.1-5 Hz bandpass filter (MATLAB 2022b). The filtered ΔF/F data were then z-scored for each recording trial.

### ΔF/F bootstrapping

The fluorescence data was bootstrapped by randomly selecting 470 frames from the entire dataset, for each CCF region, and calculating the average fluorescence value. This sample size matches the number of correct laps across all imaging trials. Using 1000 replicates, a 95% confidence interval was calculated by finding the mean randomized fluorescence ± 2 SDs. Any mean fluorescence value of the behavior-aligned data outside of this interval indicates the temporal activation produces a significantly different fluorescence pattern than random chance. All average fluorescence peaks and troughs in **Figure 4** exceeded the 95% confidence interval of the randomized data, so we can conclude these are not chance events.

### Identifying cortical activation patterns using K-means clustering

Using a clustering method adapted from a previous study^28^, the Ca^2+^ data was reduced into distinct cortical activation patterns. A k-means clustering step was implemented at three levels: 1) trial level, 2) mouse level, and 3) global level. For trial-level clustering (**Figure 4A**), the correlation coefficients between each ΔF/F frame were calculated (MATLAB *corrcoef* function) and the resulting correlation values were clustered using a k-means algorithm (5000 iterations; 5 replicates).

To determine the optimal number of clusters for a particular trial, a data reconstruction algorithm was utilized to guarantee that the resulting data reduction described the majority of the variability in the real ΔF/F data. By sequentially testing increasing numbers of clusters (starting with 2 clusters), the average of all frames within each cluster was calculated, and the resulting images represented the spatial weighting matrix. The temporal weighting matrix was calculated by multiplying the 2-dimensional ΔF/F stack with the pseudoinverse of the spatial weighting matrix (**Figure 4A**). This resulted in a time course for each cluster representing how much it contributes to each ΔF/F frame. Data reconstruction was completed by multiplying the temporal component for each frame with the spatial component. A correlation between each real ΔF/F frame and the reconstructed frame was calculated, and the average across all frames as used to determine if the clusters sufficiently described the real ΔF/F data. When the average correlation between real and reconstructed frames reached a threshold of 0.85, successive testing of cluster numbers was aborted, and that number of clusters was used for the within-trial k-means analysis. Occasionally, the reconstruction correlation with the real ΔF/F data plateaued before reaching the 0.85 threshold. In this case, the optimal cluster number was determined as the cluster number when the correlation changed less than 3% over the last 5 iterations. Using the output from the trial-level clustering, a second k-means clustering step was completed to classify a set of clusters for all trials within individual animals. This second step followed the analysis outlined for the trial-based clustering, except the optimal number of clusters was determined by the average number of clusters for the particular animal. In this clustering step, the k-means parameters were set to 10^6^ iterations and 100 replicates.

The final k-means clustering step combined all the output from the within-mouse clustering to define a universal catalog of clusters that we refer to as cortical activation states. Due to varying fields of view for each animal, the Allen Brain Atlas Common Coordinate Framework (CCF)^62^, was applied to a representative brain image for each mouse. The average pixel value within each CCF region was calculated from each output image of the mouse-level clustering. Reducing image size to the 66 average atlas regions made it possible to compare average cluster images across different mice by ensuring all regions were anatomically consistent. The correlation of each reduced image was calculated and clustered using k-means with 10^6^ iterations and 500 replicates as parameters. The optimal number of states was determined by a t-distance calculation that maximizes the within cluster correlation and minimizes the across cluster correlation^28,63^.

### Cortical state bootstrapping

To quantify significant two state transitions, random 90-frame segments of the cortical state time course were selected from the datasets of either behavioral paradigm. The number of random 90-frame segments matched the number of correct laps across all imaging trials. These segments were shuffled within each mouse to disrupt any naturally occurring transitions, and the transition probabilities of randomized data were calculated. This was repeated over 1000 iterations to create a distribution of random transition probabilities for each of the 110 possible transition pairs. Any transition probability of the real, behaviorally aligned data > 2 SDs above the mean probability of randomized data was considered significant.

To assess statistical significance for the state frequency of maze regions, the chance level probabilities were determined from the frequencies of randomly shuffled data to uncouple any real patterns of state activation and location in the maze. Again, by performing 1000 iterations of the shuffling, a distribution for each state-region probability was created and significance was determined if the real probability exceeded 5 SDs of this distribution.

### Characterizing cortical state motifs

To find cortical state motifs, the state data was first smoothed using a custom script that found the most common state within a 5-frame sliding window (MATLAB 2022b). This resulted in the elimination of many single-frame transitions, while maintaining the general patterns of state activation. During the time window ± 3 s after crossing the decision point, the frequency of each 3-state permutation was calculated, and probability was determined by normalizing to the total number of sequences. Prolonged activations of individual states were reduced to a single frame to isolate the transitions between different states. For this analysis, we chose to only utilize the time window around the decision point, as this should yield the most information about the spatiotemporal patterning of corresponding cortical motifs. Since the animal’s decision has already been made when it reaches the reward spout, we chose not to analyze this time window as it would not provide information about how the decision is generated. Only sequences of 3 different states were considered motifs as it describes a complete transformation of neural activation.

The motif probabilities generated from the real data were then compared to randomized bootstrapped data to determine which were statistically significant. A 90-frame segment was randomly selected from either the alternating or left-only data to generate a matrix that is the identical size of the real behaviorally aligned data. This data was then smoothed in the same way as the real aligned data. All states within each mouse were then shuffled individually to preserve the state distribution per mouse. Motif probability of the randomized data was then calculated, and the results were obtained over 1000 iterations. Any motifs from the real, aligned data >5 SDs above the mean randomized probability were considered significant. To isolate only the most probable motifs, a minimum probability of 0.005 was applied.

### Statistical analysis

All statistical analyses were performed using Graphpad PRISM 10.2.3. State probability curves were compared using a 2-way ANOVA with Bonferroni post-hoc correction. Each paradigm was represented by a probability curve for each of the 6 subjects. If a state was not present for a particular animal, a probability of zero was used. Samples were matched between the two paradigms since the probabilities were calculated using the same animals, during the same time period with respect to behavior. Any comparisons that showed a significant main effect but lacked significant post-hoc time points or had less than 2 consecutive post-hoc points were excluded from interpretation as we determined these differences to be too minor to make any impactful claims.

## Data and Code Availability

All calcium imaging recordings, behavioral recordings, and MATLAB scripts used in data analysis are available upon request.

## Author Contributions

Conceptualization and Funding: SH, DS, RC, SK, TE

Technology development and experiments: SH, DS, SK, LZ

Data analysis: SH, DS

Interpretation and figures: SH, TE

Project supervision: SH, AN, RC, TE

Manuscript writing and editing: SH, TE, RC, AN, LP, LZ, DS, SK

## Acknowledgements

The authors and members of the Ebner lab would like to thank Lijuan Zhuo for assisting with rodent surgeries and general laboratory support during this project. We thank members of the Kodandaramaiah lab for their work in developing the mini-mScope and constructing much of the data analysis pipeline for this project. This work was funded in part by NIH grants P30 DA048742 (TJE and SK), R01 NS111028 (SK and TJE), R01AG075809 (TJE), and RF1 NS126044 (SK and TJE).

---

**Supplemental figure 1:**
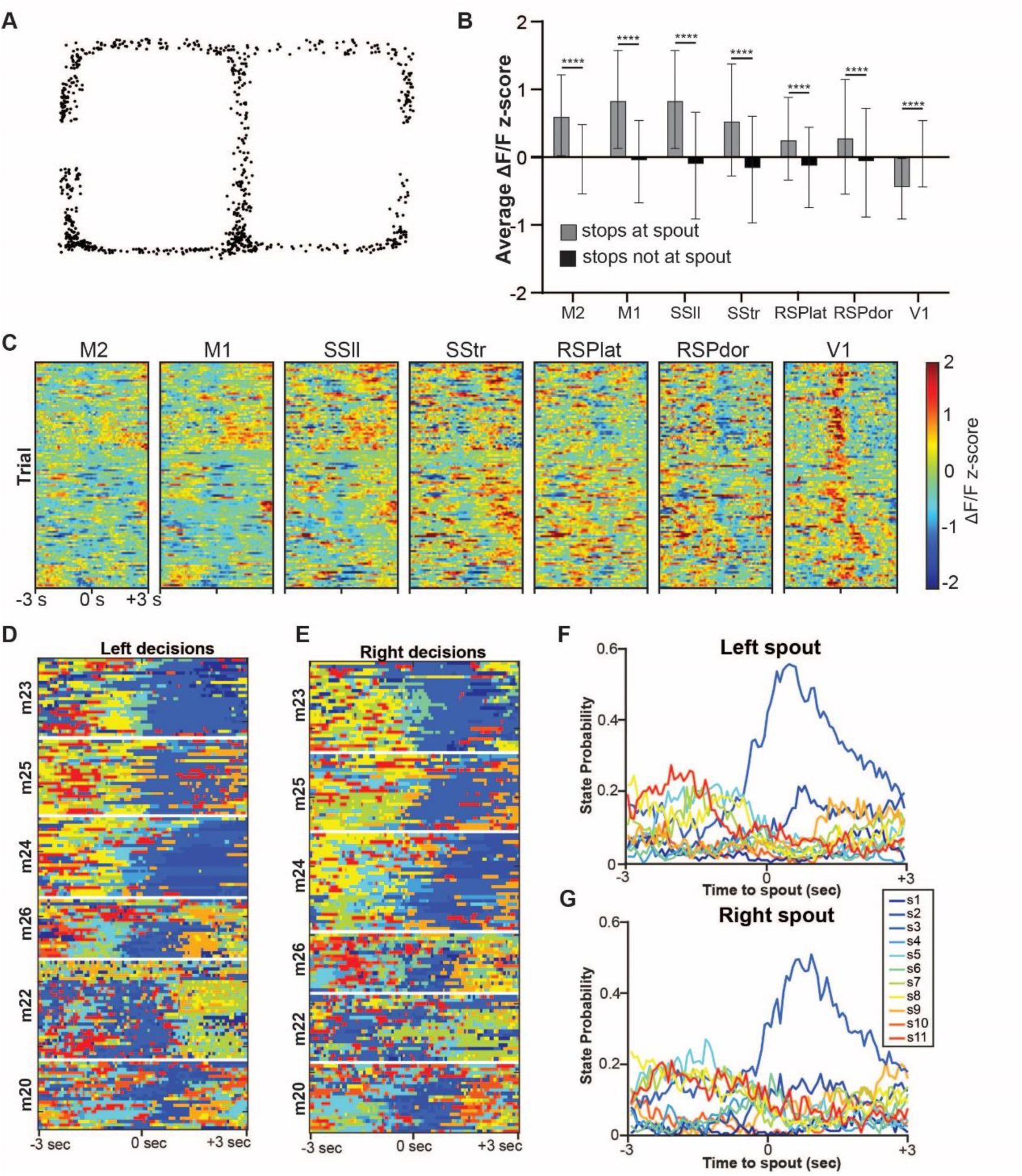
Confounding variable verification. **A.** Mouse head position at the start of every reduced locomotion period (<0.05 m/s) while outside of the two spout regions. **B.** Average ΔF/F z-scores for each of the 7 CCF regions of interest during reduced locomotion periods in the spout regions (gray) and outside of the spout regions (black). Data are presented as mean ± SD. Significant comparisons indicated with (*). **C**. ΔF/F z-score time course of each CCF region of interest over a 6 second time window centered on the moment the animal crossed the center IR beam, just before the decision point, during no-tone alternating trials. **D**. State time course over a 6 second time window centered on the moment the animal interacts with the spout during correct trials, separated by left and right-hand decisions. **F-G**. Average probability of each state at the left spout (**F**), and right spout (**G**), during the same 6 second time window as D-E.

**Supplemental figure 2:**
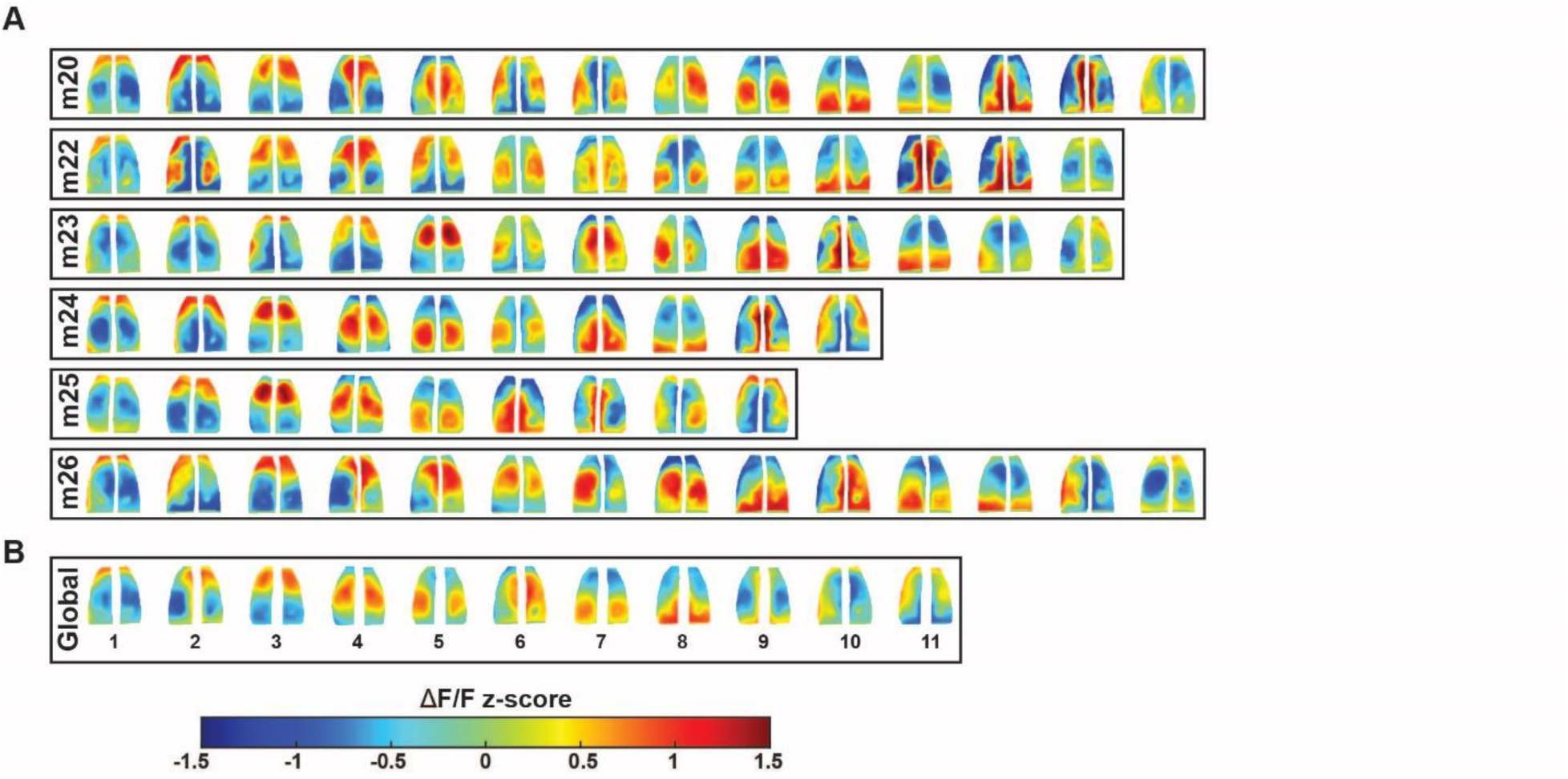
K-means clustering results. **A.** Average images for each mouse-level cluster for each of the 6 subjects. **B.** Global cortical state images from across-mouse clustering. Color scale applies to both A and B.

**Supplemental figure 3:**
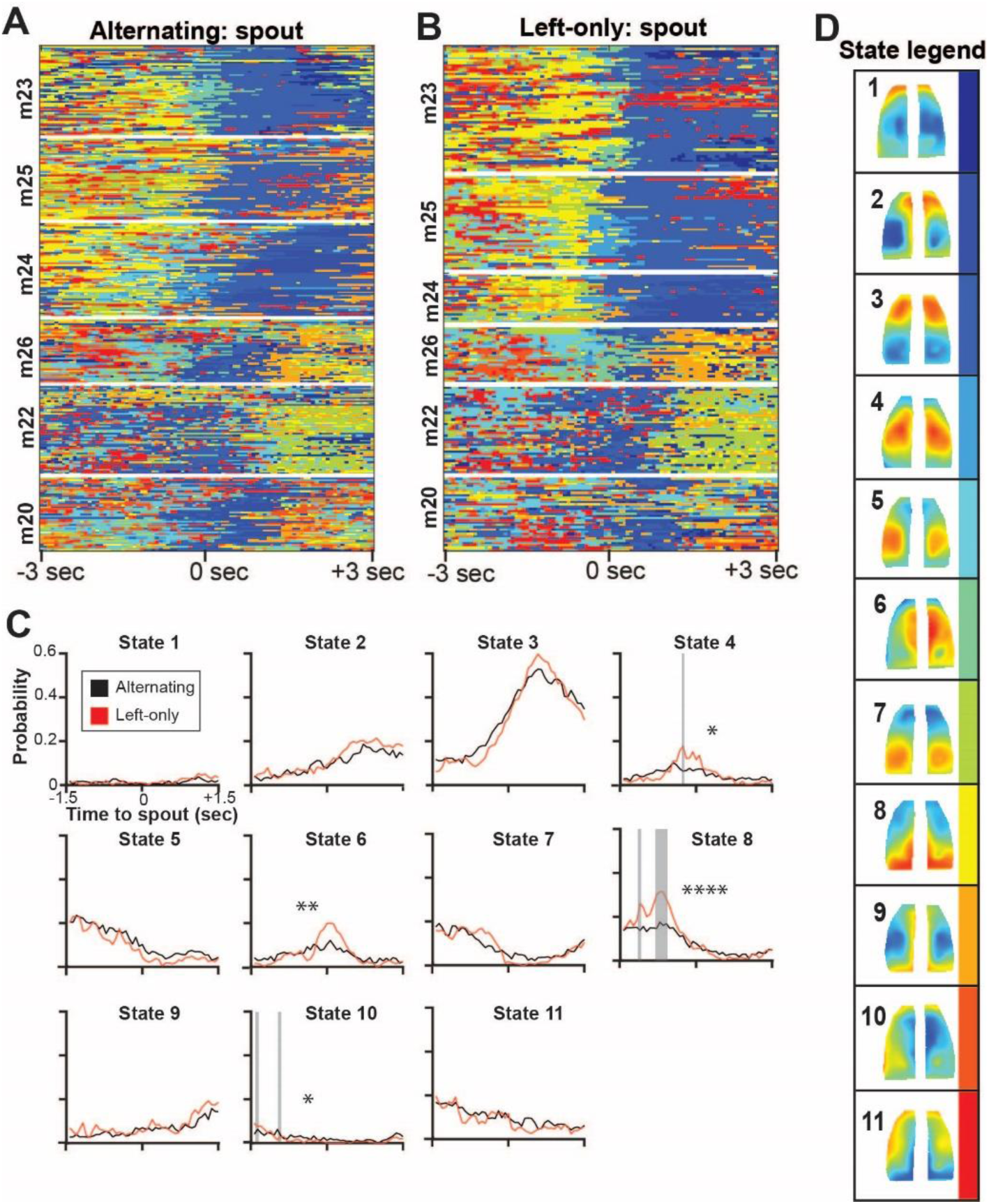
Cortical activation states aligned to the spout. **A-B.** Cortical activation state time course for all 470 correct trials during a 6 second time window centered on the spout interactions for the alternating (**A**) and left-only (**B**) paradigms. **C.** Average state probabilities for each of the 11 cortical activation states during alternating (black lines) and left-only (red lines). Significant post-hoc time points indicated with gray shaded areas. 2-way ANOVA main effect indicated with (*). **D.** Cortical activation state key.

**Supplemental figure 4:**
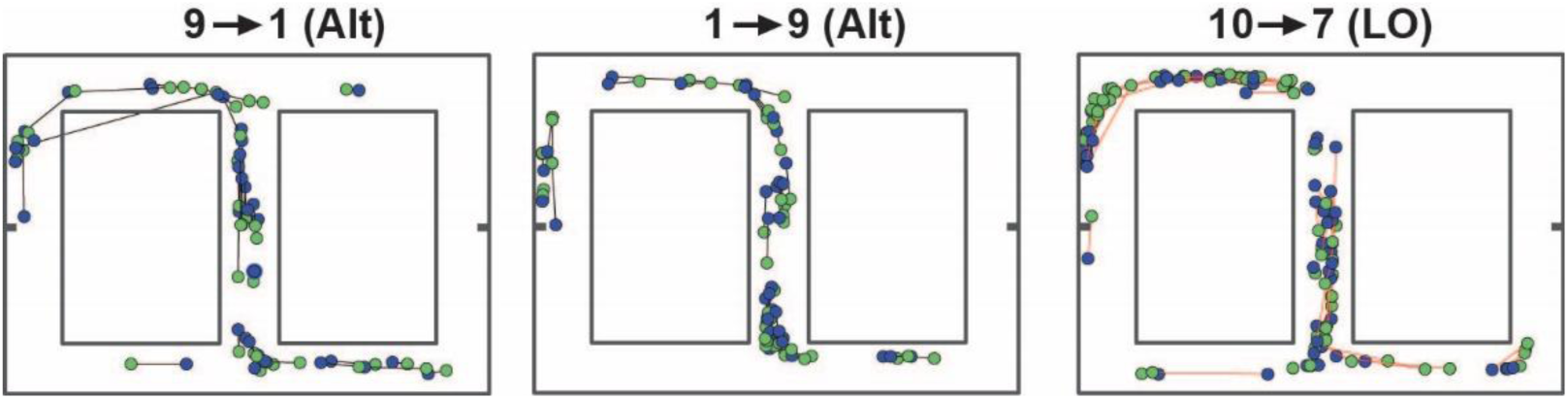
Mouse locations during significant 2-state transitions. Mouse head position during the onset of a state transition (green dots) and offset (blue dots). Onset/offset pairs are connected with a black line during the alternating paradigm, and a red line during the left-only paradigm.

